# Exponential fluorescent amplification of individual RNAs using clampFISH probes

**DOI:** 10.1101/222794

**Authors:** Sara H. Rouhanifard, Ian A. Mellis, Margaret Dunagin, Sareh Bayatpour, Orsolya Symmons, Allison Cote, Arjun Raj

**Affiliations:** Department of Bioengineering, University of Pennsylvania, Philadelphia PA; Genomics and Computational Biology Group, Perelman School of Medicine, University of Pennsylvania, Philadelphia PA; Department of Genetics, Perelman School of Medicine, University of Pennsylvania, Philadelphia PA

## Abstract

Non-enzymatic, high-gain signal amplification methods with single-cell, single-molecule resolution are in great need. We present click-amplifying FISH (clampFISH) for the fluorescent detection of RNA that combines the specificity of oligonucleotides with bioorthogonal click chemistry in order to achieve high specificity and extremely high-gain (>400x) signal amplification. We show that clampFISH signal enables detection with low magnification microscopy and separation of cells by RNA levels via flow cytometry. Additionally, we show that the modular design of clampFISH probes enables multiplexing, that the locking mechanism prevents probe detachment in expansion microscopy, and that clampFISH works in tissue samples.

## Main text

Single molecule RNA fluorescence *in situ* hybridization (RNA FISH), which enables the direct detection of individual RNA molecules^1,2,3^, has emerged as a powerful technique for measuring both RNA abundance and localization in single cells. Yet, while single molecule RNA FISH is simple and robust, the total signal generated by single molecule RNA FISH probes is low, thus requiring high-powered microscopy for detection. This keeps throughput relatively low and precludes the use of downstream detection methods such as flow cytometry. As such, amplification methods for single molecule RNA FISH with high efficiency, specificity and gain could enable a host of new applications.

A number of different signal amplification techniques are available, but each suffers from particular limitations. Approaches such as tyramide signal amplification (TSA)^4^, or enzyme ligated fluorescence (ELF)^5^ utilize enzymes to catalyze the deposition of fluorescent substrates near the probes. Alternatively, enzymes can ligate oligonucleotides to form a circular probe then catalyze a “rolling circle” nucleic acid amplification to generate a repeating sequence that can subsequently detected using fluorescent oligonucleotides^6–8^. These methods can lead to large signal gain, but must overcome the limited accessibility of (sometimes multiple) bulky enzymes through the fixed cellular environment to the target molecule. For example, the DNA ligases frequently used to circularize padlock probes are often quite inefficient^9^, contributing to inefficient amplification^10^. Meanwhile, there are a number of non-enzymatic amplification schemes, most notably the hybridization chain reaction^11–13^ and branched DNA^14–16^ techniques. These methods rely only on hybridization to amplify signal by creating larger DNA scaffolds to which fluorescent probes can attach, and so often have limited amplification potential^17^and the level of multiplexing can be limiting. Thus, our goal was to create a non-enzymatic, exponential amplification scheme with high sensitivity (detection efficiency), very high gain (signal amplification), and specificity (low background).

We first designed probes that would bind with high specificity and sensitivity; i.e., that could allow the probes to survive repeated liquid handling in conditions stringent enough to limit nonspecific binding and thus prevent spurious amplification. Padlock probes are a class of circular DNA probes that have these properties: they bind to the target region of complementarity via the 5’ and 3’ ends of the probe, with the intervening sequence not hybridized to the target in a “C” configuration.^18^ Conventionally, the ends are then connected using a DNA or RNA^19^ ligase. This connection, in combination with the DNA:RNA double helix formed upon hybridization, result in a molecule that is physically wrapped around the target strand (**Figure 1a**). We wished to retain the benefits of padlock probes without the need for this enzymatic ligation; therefore, we designed padlock-style probes with terminal alkyne and azide moieties at the 5’ and 3’ ends (click-amplifying FISH (clampFISH) probes; **Figure 1a, Supplementary Figure 1**). When the clampFISH probe hybridizes to the target RNA, the DNA:RNA hybrid brings the two moieties together in physical space. We then used a click chemistry strategy (copper(I)-catalyzed azide-alkyne cycloaddition, CuAAC^20^) to covalently link the 5’-alkyne and 3’-azide ends of the probe using small molecules rather than enzymes, wrapping around the target RNA (**Figure 1a**).

**Figure 1.**
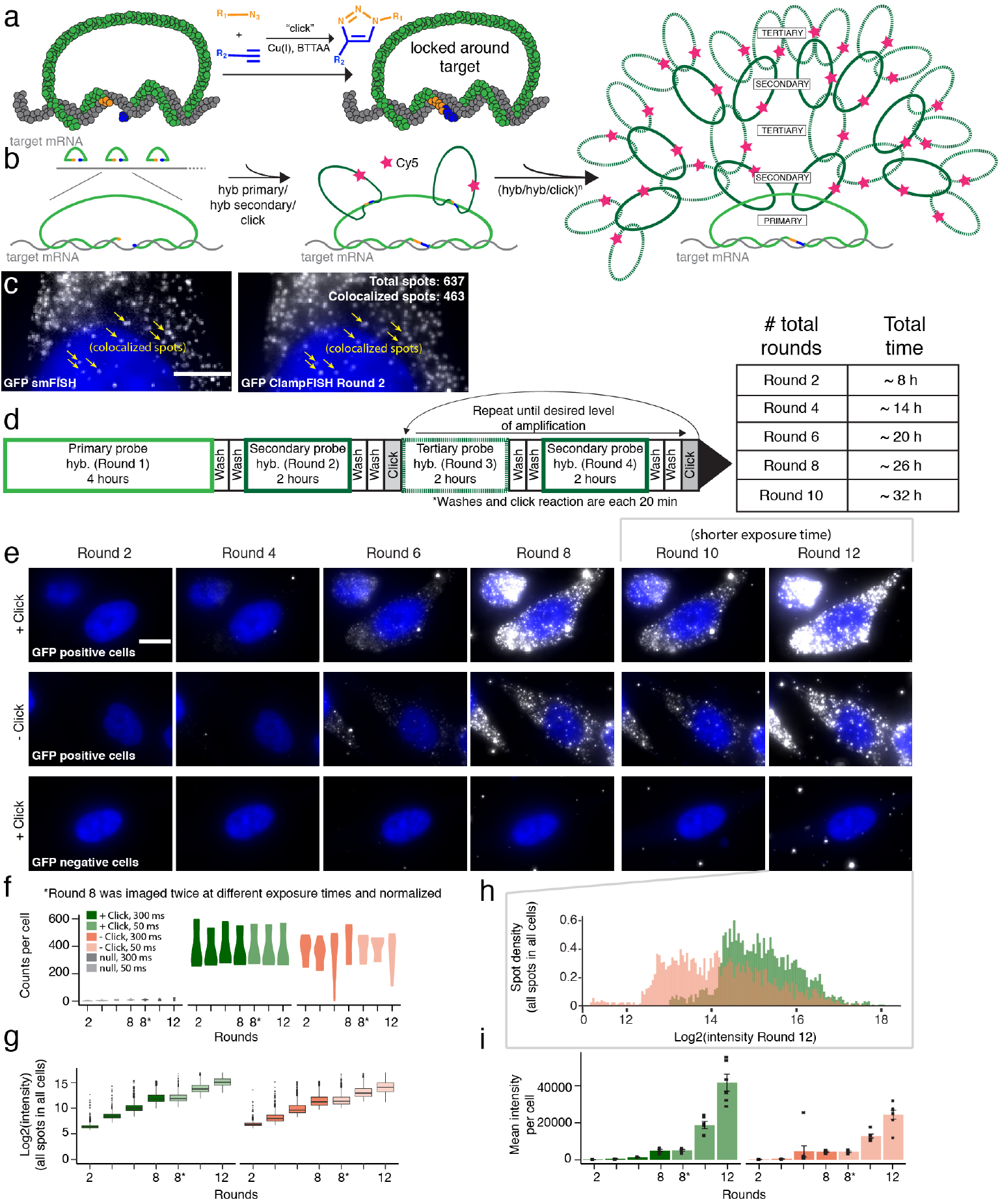
Design and validation of clampFISH technology (data shown is representative of 5 biological replicates). (**a**) clampFISH probes topologically wrap around target (left) and can be ligated to connect the 5’ and 3’ ends of the probe using CuAAC (right). (**b**) Workflow for clampFISH: multiple, primary clampFISH probes bind to the target of interest. Secondary clampFISH probes bind 2:1 to each primary clampFISH probe, and tertiary clampFISH probes bind 2:1 to each secondary clampFISH probe. In a subsequent round, the secondary probes again bind 2:1 to the tertiary probes and so on, thus providing exponential amplification. (**c**) Colocalization of GFP mRNA single molecule RNA FISH (left) with GFP mRNA clampFISH round 2 (right; scale bar = 5 μm) (**d**) Timing and order of clampFISH amplification steps. (**e**) GFP mRNA clampFISH signal on WM983b-GFP cells across 12 rounds of amplification in the presence of click ligation (top) compared to GFP mRNA clampFISH signal in the absence of click ligation (middle). Single cell tracking of the same cell line without GFP mRNA expression across rounds to assess background signal (below; scale bar = 10 μm). Images are representative single-cells selected from 3 biological replicates. (**f**) mRNA counts per cells across rounds. (**g**) Log2(intensity) of click vs. no click samples across rounds. Graphs are representative of 3 biological replicates. (**h**) Density of the log2(intensity) of all spots detected at round 12 in click vs. no click samples. (i) Mean fluorescence intensity of GFP mRNA clampFISH signal per cell on WM983b-GFP cells across 12 rounds of clampFISH. Graphs are representative of 3 biological replicates.

To achieve exponential amplification, we first designed a series of primary clampFISH probes to target the RNA sequence of interest. The backbone of each primary clampFISH probe contains two “landing pads” for a set of secondary, fluorescent, clampFISH probes. To these secondary probes, we hybridized a set of tertiary probes that again bound in a 2:1 ratio. In a subsequent round, the secondary probes again bind 2:1 to the tertiary probes and so on, thereby in principle doubling the signal in each round (**Figure 1b**). The resulting probes bind efficiently to the target, as evidenced by the colocalization of clampFISH probes with single molecule RNA FISH probes targeting the same RNA (**Figure 1c, Supplementary Figure 2**). We have observed that the non-colocalizing spots often correspond to faint smFISH spots that are not picked up by our thresholding software. Given the absence of spots in our genetic negative controls, this suggests that these faint spots may be true positives. The number of amplification rounds may be adjusted based on the desired degree of amplification required for the particular application (**Figure 1d**).

To demonstrate exponential amplification using clampFISH probes, we first targeted and amplified a GFP mRNA in a human melanoma cell line (WM983b) stably expressing GFP^21^ using 10 primary clampFISH probes (**Figure 1e**). We used stringent hybridization conditions – specifically, a higher concentration of formamide than is traditionally used for single molecule RNA FISH –to limit nonspecific probe binding while still allowing for specific binding (**Supplementary Figure 3**). As the number of rounds progressed, the average number of spots per cell remained constant (for example, at round 2 we detected a mean of 399 spots per cell ± 62 SEM and at round 10 we detected 401 spots per cell ± 36 SEM; **Figure 1f**), while the intensity of the signal as measured by fluorescence microscopy increased. At round 12, the mean signal per spot was 446-fold higher than in round 2 (**Figure 1i**). We observed a 3.387-fold increase (geometric average with a standard deviation interval of 2.55, 4.50) for every two rounds of amplification (for a 1.69-fold increase per round; **Figure 1g-i**). We also observed that, although the fold-amplification decreases slightly at later rounds, we estimate that the saturation point would be reached around round 20 (**Supplementary Figure 4**).

To assess whether the click reaction aided in the amplification process as hypothesized, we performed the same experiment in the absence of the click-ligation of the clampFISH probes. Although the number of spots detected per cell were similar (393 mean spots per cell in the clicked samples vs. 381 mean spots per cell in the non-clicked samples), we observed lower mean signal intensity (26,076 AU ± 496 SEM for non-clicked vs. 44,450 AU ± 630 standard error of the mean for clicked samples at round 12) as well as a lack of uniformity in spot intensity (coefficient of variance at round 12 for non-clicked cells = 0.94 ± 0.013 SE vs. 0.69 ±0.01 SE for clicked cells), demonstrating that the click reaction facilitated a more uniform and higher gain amplification of primary clampFISH signal (**Figure 1h, Supplementary Figure 5**). In round 12, the interquartile range of spot intensities in the clicked condition is bounded by [23879.5, 56099.7] (25th, 75th percentile) converted fluorescent units, while in the unclicked condition it is bounded by [9938.9, 32265.7] (25th, 75th percentile) converted fluorescent units (**Figure 1h**). (Although the signal was more uniform when click chemistry was used, the spread in intensity is still large enough that it may be difficult to quantitatively estimate transcript abundance in crowded environments where it is difficult to explicitly separate spots.)

To demonstrate signal specificity, we performed the same clampFISH detection and amplification on the parental cell line that did not have the GFP gene. We detected very few false-positive spots in these cells (mean of 9.77 spots per cell ± 1.45 SEM), showing that the signals were specific to the target (**Figure 1e**). While this is number is relatively low, it may potentially interfere with the detection of RNAs with low numbers of spots.

Owing to its relatively low signal intensity, single molecule RNA FISH typically requires using a microscope equipped with a high numerical aperture objective, typically requiring oil immersion. For many applications, a low magnification air objective is preferable, both for increased throughput and for simplicity of sample handling. We reasoned that the increased RNA FISH signals that clampFISH provided had the potential to make RNA FISH signals detectable by low magnification microscopy (**Figure 2a**). To test this, we mixed 20% WM983b cells stably expressing GFP with 80% WM983b cells and probed for GFP mRNA using clampFISH probes at round 6–this was the minimum number of rounds needed to clearly discern signal at the lowest magnification for this particular target. Using clampFISH, the positive cells were clearly discernible at both 20X and 10X magnification, while the conventional single molecule RNA FISH signal was not (**Figure 2a**). At lower magnification, high density mRNA spots become difficult to count, making the assay more qualitative; however, for RNAs of low abundance it has the potential to be quantitative.

**Figure 2.**
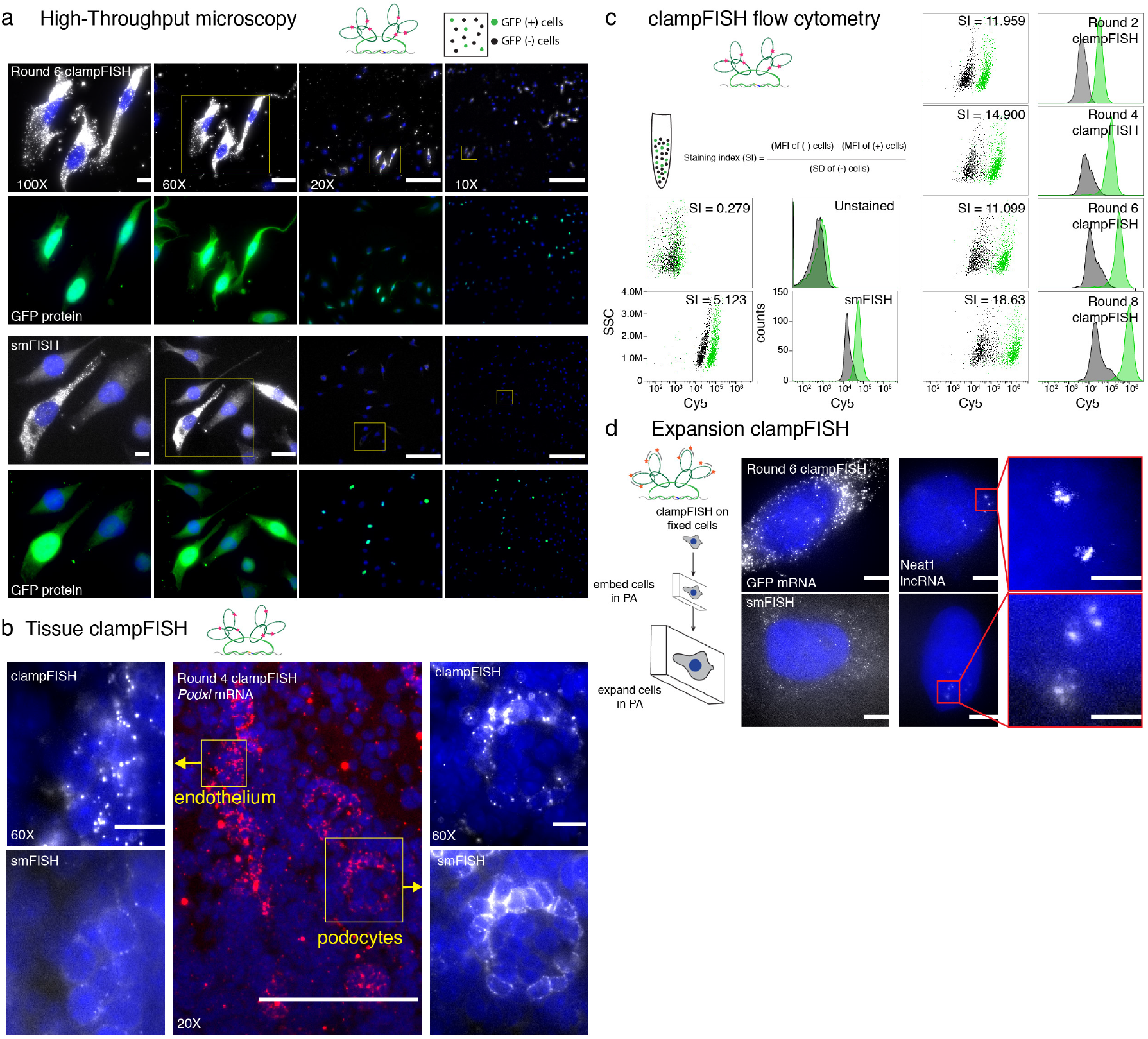
Applications of clampFISH amplification of RNA (images are contrasted independently). (**a**) 20% WM983b cells stably expressing GFP mixed with 80% WM983b cells and probed for GFP mRNA using clampFISH probes (top). Imaging was performed using 0.3 NA 10X, 0.5 NA 20X, 1.4 NA 60X and 1.4 NA 100X magnification objectives in the same positions and compared to single molecule RNA FISH of GFP mRNA using the same fluorophore (bottom; representative images of 2 biological replicates; each image is contrasted independently; scale bars are 10 μm for 100X and 60X images, 5 μm for 20X images and 2.5 μm for 10X images). Each image has a corresponding image showing GFP mRNA signal colocalizing with GFP protein. (**b**) (center) 20X image of fixed-frozen 5 μm 4do mouse kidney section stained with round 4 clampFISH probes targeting *Podxl*. (left) 60X image of mouth endothelium by round 4 clampFISH and by single molecule RNA FISH. (right) 60X image of podocyte by round 4 clampFISH and by single molecule RNA FISH (representative images shown of 2 biological replicates). (**c**) We applied clampFISH to a mixed population of MDA-MB 231 cells with and without GFP expression and analyzed the separation by flow cytometry across 8 rounds of amplification. Cells were gated on GFP expression and are displayed in green. (**d**) (top) Fluorescent micrographs of round 6 clampFISH targeting GFP mRNA and Neat1 lncRNA in cultured WM983b-GFP cells (bottom) fluorescent micrographs of single molecule RNA FISH targeting GFP mRNA and Neat1 lncRNA in cultured WM983b-GFP cells using the same dye (images representative of 2 biological replicates; scale bars are 20 μm for the images, and 5 μm for the inlay).

Primary tissue samples typically suffer from high background levels that contribute to a low signal-to-noise ratio using single molecule RNA FISH, and therefore require high magnification microscopy to discern positive signal from background. However, at high magnification, large structural features of the tissue are often difficult to discern, and tiled image scanning is relatively slow. To reduce the magnification and increase the visible area while still viewing individual RNAs, we applied 4-rounds of clampFISH to 4 day old C57BL/6J mouse kidney samples and probed for *Podxl* mRNA, a gene that is highly expressed in podocytes^22^, and observed specific expression of clampFISH signal in the appropriate regions (**Figure 2b, Supplementary Figure 6**). We chose to stop after 4 rounds because this was the minimum number of rounds to discern signal using 20X magnification. Interestingly, clampFISH further revealed that *Podxl* also expressed in the kidney endothelium, a signal that was only faintly visible by single molecule RNA FISH, but was clearly detected by clampFISH at low magnification (**Figure 2b**). This is consistent with previous findings that *Podxl* is expressed at low levels in the kidney endothelium^23^ and highlights the utility of clampFISH for detection of low abundance transcripts in tissue.

Another important application that clampFISH enables is flow cytometry-based measurement of RNA expression, an application for which single molecule RNA FISH typically does not produce enough signal^24,25^. We applied clampFISH to a mixed population of MDA-MB 231 cells with and without GFP expression and analyzed the cells by flow cytometry (**Figure 2c, Supplementary Figure 7**), using GFP fluorescence as an independent measure of the specificity of clampFISH signal. We observed separation of GFP positive cells by clampFISH signal with as few as 2 rounds of amplification, and observed a 2.447-fold increase in fluorescence intensity in the GFP positive population with every 2 rounds of amplification thereafter (geometric mean of fold change across rounds; **Figure 2c, Supplementary Figure 7-8**). Notably, we observed a decreasing fold-change as we moved through the rounds (3.435-fold from rounds 2-4, 2.589-fold from rounds 4-6, and 1.648-fold from rounds 6-8).

Amplification of RNA signal can also be used in combination with a newly developed expansion microscopy technique that achieves super-resolution microscopy via the physical expansion of cells embedded in polymeric hydrogels^26,27^. When combined with single molecule RNA FISH, expansion microscopy can resolve the fine structure of RNAs that are in close proximity to one another; however, the physical expansion of cells results in reduced signal intensities, at least partially due to probes dissociating under the low salt conditions required to obtain high levels of hydrogel expansion. We reasoned that the locking property of clampFISH probes would allow us to maintain signal intensity in the face of these expansion conditions. We thus performed clampFISH on GFP mRNA to round 6 followed by expansion and observed high signal intensity on all spots when the click reaction was performed, but with little signal when click was not performed (**Figure 2d, Supplementary Figure 9**). We also applied clampFISH to amplify *NEAT1* to round 6, a nuclearly retained long non-coding RNA, and observed higher signal intensity than with single molecule RNA FISH (**Figure 2d, Supplementary Figure 9**). This also suggests that clampFISH probes are accessible to the nucleus, which can be a problem with other amplification schemes^16^. Interestingly, we also observed nuclear localization of the GFP using clampFISH probes (**Figure 1c and e**) with the exception of transcription sites. Upon further analysis, we determined that clampFISH probes can enter the nucleus but may have difficulty accessing transcription sites (data not shown), possibly due to crowding from RNA secondary structure or nearby proteins.

A key design goal for *in situ* hybridization methods is the ability to detect multiple RNA targets simultaneously. Multiplexing with clampFISH is in principle straightforward because of the modular design of the probes. The backbone sequence of the clampFISH probes can easily be changed, allowing one to use multiple independent amplifiers simultaneously. Many transcripts may be amplified simultaneously with unique backbone sequences that are not labeled with a fluorophore, and the subsequent loop-dendrimer structure can be probed with fluorescently labeled secondary fluorescent oligonucleotides that can be easily removed and re-hybridized. As a proof-of-concept, we selected 3 RNA targets with distinct expression patterns in HeLa cells: *NEAT1*, which is found in nuclear paraspeckles of most cells; *LMNA*, which is found in the cytoplasm of all cells, and *HIST1H4E*, which expresses only in the subpopulation of cells that are in S phase. We amplified these with unique sets of non-fluorescent clampFISH probes to 7 rounds, then probed the terminal backbones with single molecule RNA FISH probes, each labeled with different fluorophores (**Figure 3a, b**). We were able to visualize signals from the three different probe sets, even using low magnification microscopy. Unexpectedly, we observed that *LMNA* was present in the transcription sites and therefore colocalized with the *NEAT1* signal. To confirm that this was not bleedthrough, we imaged each probe, amplified to 7 rounds in every channel using the same exposure times. We saw no bleedthrough between fluorescence channels (**Supplementary Figure 10**). Additionally, we confirmed that the colocalization was not due to cross-hybridization of the terminating fluorescent oligonucleotides (data not shown).

**Figure 3.**
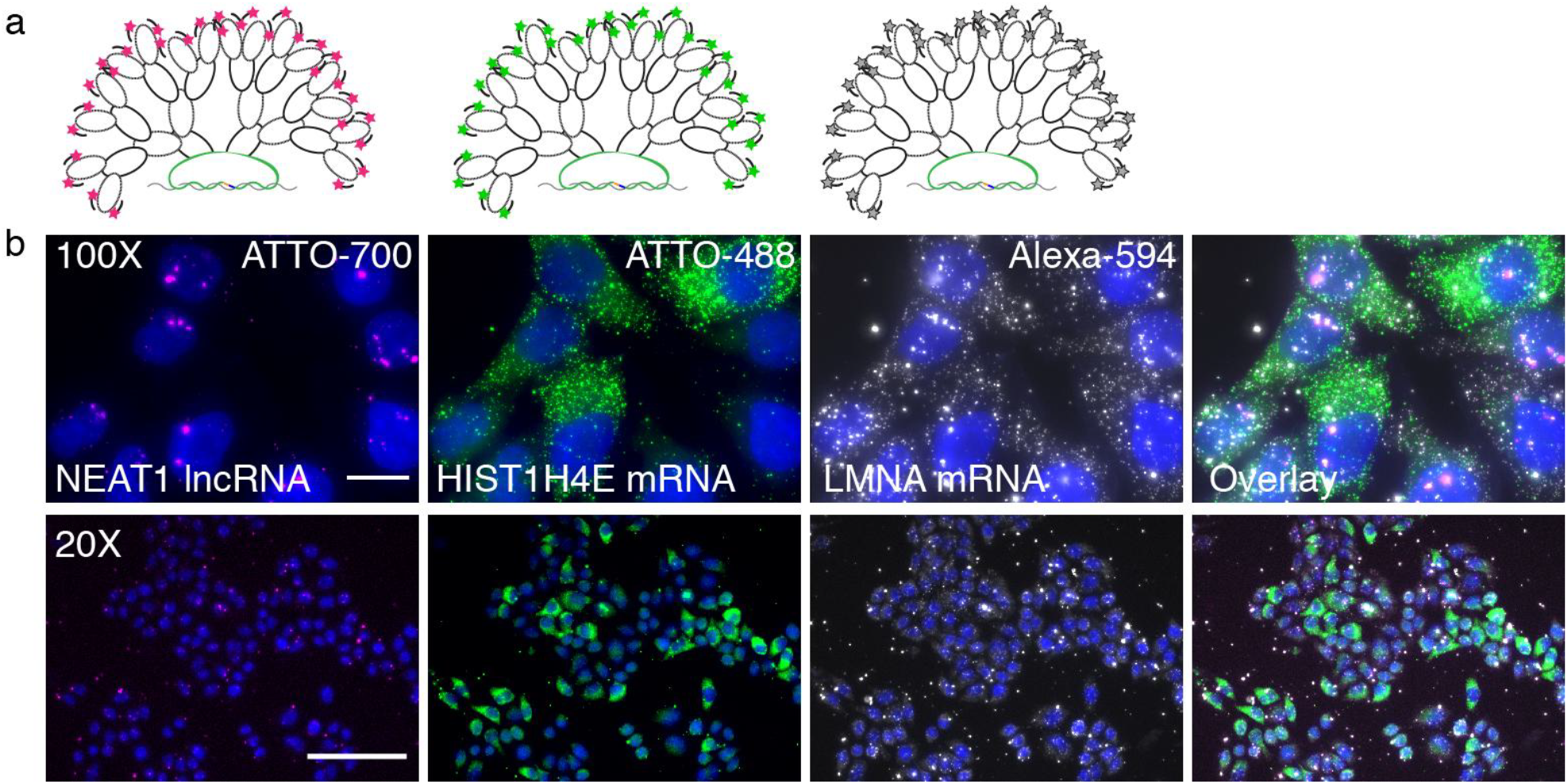
Multiplexing 3 RNA targets on HeLa cells. (**a**) Schematic diagram of probe hybridization scheme. Unique clampFISH probe sets are designed for each target, and probed at the final round with a single molecule RNA FISH probe labeled with a unique fluorophore (represented with □). (**b**) Fluorescent micrographs of individual probe channels: (from left) *NEAT1* lncRNA labeled with ATTO700, *HIST1H4E* mRNA labeled with ATTO 488, and *LMNA* mRNA labeled with Alexa 594 and an overlay on the far right. (top) 100X magnification with 20 μm scale bars, (bottom) 20X magnification with 20 μm scale bars. Images are representative of 2 biological replicates.

We present here are new scheme for the fluorescent, non-enzymatic amplification of fluorescent RNA signal *in situ*. This method is different from other non-enzymatic, hybridization-based schemes because it directly links the probe to the target RNA whereas other systems are susceptible to probe detachment during washes. In a direct comparison with commercially available systems, we observed that other methods’ maximum fluorescence intensity amplification is comparable to clampFISH at 6 rounds (**Supplementary Figure 11**), however, the fluorescence intensity of clampFISH far surpasses other methods beyond round 6; thus, the clampFISH amplification system can enable assays that require extremely high signal gain, especially flow cytometry (**Figure 2c**) and high throughput microscopy of targets with lower expression levels (**Figure 3b**). For instance, we were able to detect *HIST1H4E* RNA with low-power microscopy even though it is typically expressed at levels of only around 200 molecules per cell^21^ (**Figure 3b**). ClampFISH also may exhibit lower levels of background as compared to HCR (**Supplementary Figure 11**). Additional benefits to clampFISH include tunable, exponential amplification of fluorescence intensity (**Figure 1**) in addition to modular probe design for simplified and expanded multiplexing capabilities. The resulting method, clampFISH, enables the probing and visualization of individual RNAs on cells and tissues using low powered microscopy and is compatible with expansion microscopy. Additionally, clampFISH amplification may be used to separate cells based on their RNA expression using flow cytometry. Interestingly, the combination of probe hybridization and click chemistry moieties on the ends of the primary clampFISH probes behave as a proximity ligation wherein the click reaction will occur if and only if the two arms are hybridized adjacent to each other (**Supplementary Figure 12**). These data suggest that clampFISH may in future find uses in specifically probing other difficult-to-image RNA subsets such as splicing junctions, short alternatively spliced variants, or edited RNAs.

## Materials and Methods

### Cell culture

We cultured WM983b cells and WM983b-GFP-NLS cells (a human metastatic melanoma cell line from the lab of Meenhard Herlyn) in tumor specialized media containing 2% FBS. The WM983b-GFP-NLS contains EGFP fused to a nuclear localization signal driven by a cytomegalovirus promoter that we stably transfected into the parental cell line.

### Clamp probe design and synthesis

Clamp probes are 150 nt long (15mer left RNA binding arm, 10 nt left adapter, 100mer backbone, 10 nt right adapter, 15mer right RNA binding arm). RNAs are targeted by probe sets containing one or more Clamp probes, each targeting a 30 nt region of RNA (2 adjacent 15mer binding arms). We chose binding regions with approximately 40% GC content as well as minimal repetitive regions using our probe design served (source code available here: https://flintbox.com/public/project/50547/) and instructions for use are available in the **supplementary methods**. We designed backbones so as to minimize secondary structure (using mFold; http://unafold.rna.albany.edu/?q=mfold). We ordered modified DNA oligonucleotides from Integrated DNA Technologies (IDT) as standard DNA oligonucleotides with modifications (5′-phosphate on the backbone, 3′-azide and 5′-phosphate for the right arm and 5′-hexynyl for the left arm). Strands were resuspended in nuclease free water, at a working stock concentration of 400 μM. The left arm (30 μM), backbone (20 μM) and right arm (30 μM) are brought together using adapter probes (30 μM each) and heated to 70C for 3 min prior to being enzymatically ligated using 600 U of T7 DNA ligase (New England Biolabs) for a minimum of 1 hour at room temperature. Following ligation, the probes were purified using Monarch purification columns (New England Biolabs) and eluted in 4X the starting volume to make the working dilution. For a schematic protocol and probe sequences, see **Supplementary figure 1 and Supplementary table 1**.

### ClampFISH procedure on cultured cells

We grew cells on glass coverslides until ~70% confluent. We washed the cells twice with 1X PBS, then fixed for 10 minutes with 4% formaldehyde/1X PBS at room temperature. We aspirated off the formaldehyde, and rinsed twice with 1X PBS prior to adding 70% ethanol for storage at 4°C. We incubated our cells for at least 4 hours at 37°C in hybridization buffer (10% dextran sulfate, 2X SSC, 20% formamide) and 1 μl of the working dilution of the primary ClampFISH probe. We performed two washes in wash buffer (2X SSC, 10% formamide), each consisting of a 30-min incubation at 37°C. We then incubated the cells for at least 2 hours at 37°C in hybridization buffer (10% dextran sulfate, 2X SSC, 20% formamide) and 1 μl of the working dilution of the secondary ClampFISH probe and repeated the washes. After the second wash, we performed the ‘click’ reaction. A solution containing 75 μM CuSO_4_·5H2O premixed with 150 μM BTTAA ligand^20^ (Jena Biosciences) and 2.5 mM sodium ascorbate (made fresh and added to solution immediately before use; Sigma) in 2X SSC was added to the samples, and these were then incubated for 30 min at 37°C. The samples were then rinsed briefly with wash buffer, then we continued cycling this protocol, alternating between secondary and tertiary ClampFISH probes until reaching the desired level of amplification. After the final wash, we rinsed once with 2X SCC/DAPI and once with anti-fade buffer (10 mM Tris (pH 8.0), 2X SSC, 1% w/v glucose). Finally, we mounted the sample for imaging in an anti-fade buffer with catalase (Sigma) and glucose oxidase^2^ (Sigma) to prevent photobleaching.

### ClampFISH for flow cytometry

ClampFISH for flow cytometry was performed as described above however the cells were kept in suspension. Wash buffer and 2X SSC were supplemented with 0.25% Triton-X, and the clampFISH hybridization buffer was supplemented with the following blocking reagents: 1μg/μl yeast tRNA (Invitrogen), 0.02% w/v bovine serum albumin, 100ng/μl sonicated salmon sperm DNA (Agilent).

### ClampFISH for expansion microscopy

Acryloyl-X, SE (6-((acryloyl)amino)hexanoic acid, succinimidyl ester, here abbreviated AcX; Thermo-Fisher) was resuspended in anhydrous DMSO at a concentration of 10 mg/mL, aliquoted and stored frozen in a desiccated environment. Label-IT ® Amine Modifying Reagent (Mirus Bio, LLC) was resuspended in the provided Mirus Reconstitution Solution at 1mg/ml and stored frozen in a desiccated environment. To prepare LabelX, 10 μL of AcX (10 mg/mL) was reacted with 100 μL of Label-IT ® Amine Modifying Reagent (1 mg/mL) overnight at room temperature with shaking. LabelX was subsequently stored frozen (−20 °C) in a desiccated environment until use. Fixed cells were washed twice with 1× PBS and incubated with LabelX diluted to 0.002 mg/mL in MOPS buffer (20 mM MOPS pH 7.7) at 37 °C for 6 hours followed by two washes with 1× PBS.

Monomer solution (1x PBS, 2 M NaCl, 8.625% (w/w) sodium acrylate, 2.5% (w/w) acrylamide, 0.15% (w/w) N,N’-methylenebisacrylamide) was mixed, frozen in aliquots, and thawed before use. Prior to embedding, monomer solution was cooled to 4°C to prevent premature gelation. Concentrated stocks (10% w/w) of ammonium persulfate (APS) initiator and tetramethylethylenediamine (TEMED) accelerator were added to the monomer solution up to 0.2% (w/w) each. 100 μl of gel solution specimens were added to each well of a Lab Tek 8 chambered coverslip and transferred to a humidified 37° C incubator for two hours. Proteinase K (New England Biolabs) was diluted 1:100 to 8 units/mL in digestion buffer (50 mM Tris (pH:8, 1 mM EDTA, 0.5% Triton X-100, 0.8 M guanidine HCl) and applied directly to gels in at least ten times volume excess. The gels were then incubated in digestion buffer for at least 12 hours. Gels were then incubated with wash buffer (10% formamide, 2× SSC) for 2 hours at room temperature and hybridized with RNA FISH probes in hybridization buffer (10% formamide, 10% dextran sulfate, 2× SSC) overnight at 37 °C. Following hybridization, samples were washed twice with wash buffer, 30 minutes per wash, and washed 4 times with water, 1 hr per wash, for expansion. Samples were imaged in water with 0.1μg/mL DAPI.

### ClampFISH for mouse tissues

All studies were carried out under a protocol approved by the Institutional Animal Care And Use Committee at the University of Pennsylvania. Kidneys were harvested from 4 day old C57BL/6J mice. Dissected tissues were embedded in OCT, then flash frozen using liquid nitrogen. 5 μm tissue sections were cut at −20°C and mounted on charged slides. Slides were washed briefly in PBS, then immersed in 4% paraformaldehyde for 10 min at room temperature. Following fixation, the slides were transferred to 70% ethanol for permeabilization for at least 12 hours, or for long-term storage. To begin clampFISH procedure, slides were transferred to wash buffer for 3 minutes to equilibrate, then 500 μl of 8% SDS was added to the top of the flat slide for 1 minutes for tissue clearing. Samples were transferred to wash buffer, and normal clampFISH procedure was used.

### ClampFISH for multiplexing

Primary clampFISH probes for multiple targets (each with a different backbone series) are hybridized and washed at the same time. Each subsequent round is performed together using the respective secondary and tertiary probes that are colorless. After the terminal round, samples are washes with 10%formamide/2X SSC, then hybridized with RNA FISH probes in hybridization buffer (10% formamide, 10% dextran sulfate, 2× SSC) overnight at 37 °C. Following hybridization, samples were washed twice with wash buffer, 20 minutes per wash, then counterstained with Dapi nuclear stain and prepared for imaging.

### Comparison of amplification methods

GFP mRNA clampFISH was performed to round 6 according to the protocol reported above on WM983b-GFP and WM983b cells. These probes were Cy5 labeled, and signal intensity was compared to a corresponding smFISH probe set targeting GFP mRNA that was also labeled with Cy5. GFP mRNA was also detected using RNAscope technology © according to the manufacturer’s protocol (ACDbio). These probes were ATTO 647 labeled, and signal intensity was compared to a corresponding smFISH probe set targeting GFP mRNA that was also labeled with ATTO 647. GFP mRNA was also detected using HCR technology © according to the manufacturer’s protocol (Molecular Instruments). These probes were Alexa 647 labeled, and signal intensity was compared to a corresponding smFISH probe set targeting GFP mRNA that was also labeled with Alexa 647.

### Imaging

We imaged each samples on a Nikon Ti-E inverted fluorescence microscope a cooled CCD camera (Andor iKon 934). For 100× imaging, we acquired z-stacks (0.3 μm spacing between stacks) of stained cells. The filter sets we used were 31000v2 (Chroma), 41028 (Chroma), SP102v1 (Chroma),17 SP104v2 (Chroma) and SP105 (Chroma) for DAPI, Atto 488, Cy3, Atto 647N/Cy5 and Atto 700, respectively. A custom filter set was used for Alexa 594 (Omega). We varied exposure times depending on the dyes and degree of amplification used. Typically, ClampFISH imaging was done at a 300 ms exposure and single molecule RNA FISH was done at 2-3 s exposure.

### Image analysis

We first segmented and thresholded images using a custom MATLAB software suite (downloadable at https://bitbucket.org/arjunrajlaboratory/rajlabimagetools/wiki/Home). Segmentation of cells was done manually by drawing a boundary around non-overlapping cells. Unless otherwise specified, we called clampFISH and single molecule RNA FISH spots using the previously described algorithm in rajlabimagetools (https://bitbucket.org/arjunrajlaboratory/rajlabimagetools/wiki/Home). *For colocalization analysis: We performed spot colocalization analysis as previously described^28^*. Briefly, after initial spot calling, the algorithm used Gaussian fitting to refine spot localization estimates, and then a 2-stage algorithm incorporating chromatic aberration correction to identify pairs of spots colocalizing across two channels. *For expansion-FISH:* We manually segmented cells as described above. We performed spot calling using the modified expansion spot-calling processor in rajlabimagetools, as previously described.

### Statistical analysis

All experiments were performed in replicate. Error bars throughout represent standard error of the mean unless otherwise specified.

### Reproducible analyses

Scripts for all analyses presented in this paper, including all data extraction, processing, and graphing steps are freely accessible at the following url: https://www.dropbox.com/sh/b2mv4o9wmzcicqv/AABARZsKtD1TQKMseoG_LnyWa?dl=0.

Our image analysis software is available here: https://bitbucket.org/arjunrajlaboratory/rajlabimagetools/wiki/Home, changeset 6aa67c3b68c8dd5599fed681e1a21ec674464c65. All raw and processed data used to generate figures and representative images presented in this paper are available at the following url: https://www.dropbox.com/sh/b2mv4o9wmzcicqv/AABARZsKtD1TQKMseoG_LnyWa?dl=0.

## Acknowledgements

We thank F. Tuluc from the CHOP flow cytometry core facility for helpful discussions and assistance with flow cytometry. S.H.R. acknowledges support from NIH 1F32GM120929-01A1; I.A.M. acknowledges support from NIH F30 NS100595; O.S. acknowledges support from the Human Frontier Science Program LT000919/2016-L; and A.R. from NIH 4DN U01 HL129998, NIH Center for Photogenomics RM1 HG007743, the Chan Zuckerberg Initiative, NSF CAREER 1350601, NIH R33 EB019767. We also thank the many BioRxiv readers that reached out with helpful feedback and suggestions.

